# SARM1 is an essential component of neuronal Parthanatos

**DOI:** 10.1101/2025.05.14.654090

**Authors:** Tong Wu, Liya Yuan, Yo Sasaki, William Buchser, A Joseph Bloom, Aaron DiAntonio, Jeffrey Milbrandt

## Abstract

The NAD^+^ hydrolase SARM1 is the central executioner of pathological axon degeneration. SARM1 is allosterically activated by an increased NMN/NAD^+^ ratio resulting from depletion of NAD^+^ or accumulation of its precursor, NMN, typically due to loss of the labile NAD^+^ synthetase NMNAT2 following axon injury. Another NAD^+^ hydrolase, PARP1, is hyperactivated by DNA damage, triggering the Parthanatos cell death pathway. We demonstrate that multiple mechanistically-distinct DNA-damaging agents lead to SARM1 activation and axon degeneration following PARP1 activation. Remarkably, SARM1 is required for key steps downstream of PARP1 activation by DNA damage that are pathognomonic of Parthanatos, including mitochondrial depolarization, nuclear translocation of AIF (apoptosis-inducing factor), and cell death. Moreover, SARM1 mediates glutamate excitotoxicity, a clinically significant pathomechanism attributed to Parthanatos. The identification of SARM1 as an essential component of neuronal Parthanatos, a major contributor to cell death in neurodegenerative disease, greatly expands the potential clinical utility of SARM1 inhibitors.

## INTRODUCTION

The inducible NAD^+^ hydrolase SARM1 is a metabolic sensor of axonal NAD^+^ homeostasis and the central executioner of the programmed axon degeneration pathway (AxD) responsible for Wallerian degeneration^1^. In canonical AxD, injury or disease result in loss of the NAD^+^ biosynthetic enzyme NMNAT2, a consequent decrease in axonal NAD^+^, and accumulation of its precursor, NMN^2,3^. Elevation of the NMN/NAD^+^ ratio activates SARM1, which swiftly depletes the remaining NAD^+^ and causes local metabolic collapse and axon dissolution^4,5^. Because NMN and NAD^+^ compete for the same allosteric site, SARM1 can be activated by either a rise in NMN, such as occurs after NMNAT2 loss, or by a reduction of NAD^+^, such as following activation of NAD^+^ consuming enzymes^5,6^.

Loss or inhibition of SARM1 dramatically suppresses axon degeneration in many disease and neural injury models, including hereditary and chemotherapy induced neuropathy, ocular disorders, traumatic brain injury, viral infection, and neurodegenerative conditions including CMT and ALS^7–16^. In addition to its well-described role in AxD, SARM1 can also contribute to loss of neuronal soma^10–12,15,17,18^, including cell death due to direct activation of SARM1 by neurotoxins^19,20^. SARM1-associated cell death is mechanistically different from other forms of cell death, unaffected by inhibitors of apoptosis or necroptosis^21–23^. Hence, SARM1 does not act exclusively in axons, but also mediates neuron death via a currently poorly understood mechanism.

Parthanatos, another caspase-independent death pathway, is a primary contributor to neuron death in models of Parkinson’s and Alzheimer’s disease, ALS, traumatic brain injury, and stroke^24^. Poly (ADP-ribose) polymerase 1 (PARP1), a major consumer of NAD^+^, is the key initiator of Parthanatos. In cells that experience moderate DNA damage, PARP1 hydrolyzes NAD^+^ to generate poly-ADP ribose (PAR), which recruits DNA repair factors. However, excess DNA damage—such as caused by some chemotherapeutic agents or high levels of reactive oxygen species (ROS)—hyperactivates PARP1 leading to cell death via Parthanatos^25,26^.

Typically, DNA alkylating agents like Methyl-N’-Nitro-N-Nitrosoguanidine (MNNG) are used to experimentally induce Parthanatos^27,28^. A more disease-relevant mechanism of DNA damage occurs when mitochondria produce the genotoxin peroxynitrite (ONOO−) downstream of glutamate excitotoxicity^29,30^. In all cases, DNA damage-induced PARP1 hyperactivation leads to NAD^+^ depletion and PAR formation. Parthanatos entails PAR polymers translocating to the cytoplasm, mitochondrial depolarization, association of PAR with apoptosis-inducing factor (AIF), release of AIF from the mitochondria, and AIF binding to the potent nuclease MIF (Macrophage migration inhibitory factor). The AIF-MIF complex translocates to the nucleus leading to fragmentation of the genome and cell death^31^. Thus, effectuated by mitochondrial resident components, nuclear DNA damage is both the beginning and end of the Parthanatos pathway.

The mechanism of SARM1 activation by increased NMN/NAD^+^ prompted us to examine whether PARP1 hyperactivation activates SARM1. Here, we show that PARP1 activation is sufficient to activate SARM1 and that SARM1 is necessary for key events in Parthanatos. Loss of SARM1 prevents mitochondrial depolarization, AIF translocation to the nucleus, and cell death caused by mechanistically diverse DNA damaging agents. Indeed, SARM1 inhibition is as effective as PARP1 inhibition in blocking cell death in these paradigms, identifying SARM1 as an essential component of neuronal Parthanatos. Moreover, loss or inhibition of SARM1 is profoundly protective in other Parthanatos models including DNA-damage exacerbated by ALS-associated *FUS* mutations, the Parkinsonian neurotoxin MPP^+^, and glutamate excitotoxicity. Due to its broad participation in neuropathology, targeting Parthanatos is an active area of therapeutic development. Excitingly, SARM1 inhibitors are in clinical trials and may have favorable side-effect profiles since SARM1 deletion has no obvious health consequences in animals. Thus, the identification of SARM1 as a central component of Parthanatos provides an attractive therapeutic target within this consequential neuronal death pathway.

## RESULTS

### DNA damage induces SARM1-dependent axon degeneration

Topoisomerase inhibitors are commonly used chemotherapeutic agents that damage DNA to kill rapidly dividing cancer cells. A side effect of their use is chemotherapy-induced peripheral neuropathy due to axon damage. To study the mechanism of neurodegeneration following DNA damage, we treated mouse DRG neurons with various DNA-damaging agents: the Topoisomerase I inhibitor camptothecin (CPT), the Topoisomerase II inhibitor etoposide, and MNNG, a DNA alkylating agent widely used in the laboratory to induce DNA damage^32–34^. We optimized the dose of each drug, verified DNA damage by γH2AX (Supplemental Fig 1A-C), and used brightfield imaging to assess axonal integrity over time (Fig 1A). MNNG (500 μM) provoked axon degeneration within 12 hr (Fig 1A), while degeneration in response to the other DNA damaging agents began somewhat later (Fig 1B, C, Supplemental Fig 1J). As mitochondrial dysfunction is associated with DNA damage, we measured mitochondrial membrane potential using tetramethylrhodamine methyl ester (TMRM) and found that DNA damaging agents initiated a gradual mitochondrial depolarization in accord with the rate of axon degeneration (Supplemental Fig 1D-I). These results demonstrate a link between DNA damage and axon degeneration.

**Figure 1.**
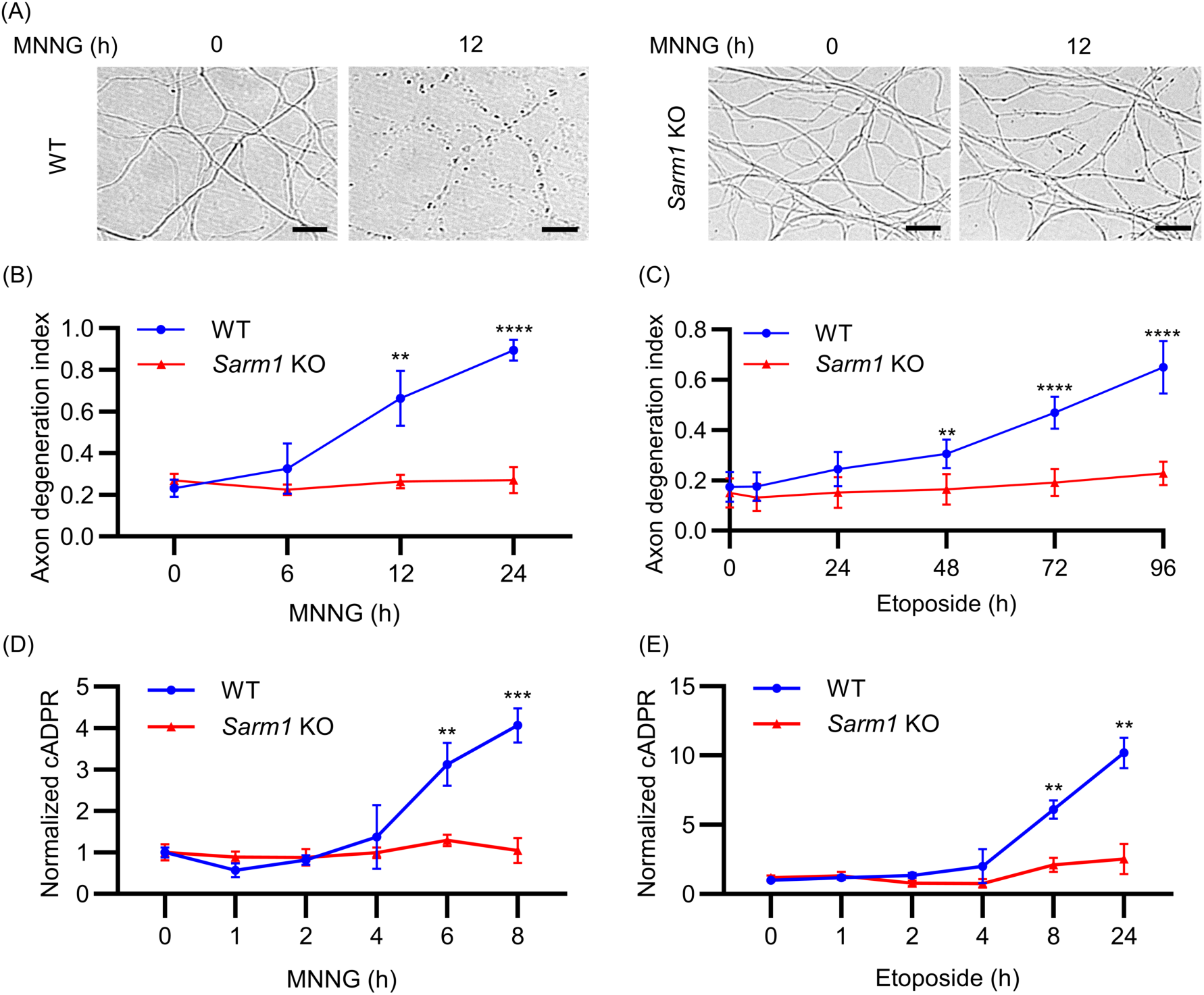
DNA damage induces SARM1-dependent neuronal degeneration. (A) WT and *Sarm1* KO DRGs were treated at DIV7. Representative images of axons from cultured WT and *Sarm1* KO DRG neurons 0 and 12 hr after treatment with 500 µM MNNG show axon degeneration. Scale bars, 25 µm. (B-C) Axon degeneration index after treatment with (B) 500 µM MNNG and (C) 300 µM etoposide. (D-E) cADPR quantified by LC-MS/MS after treatment with (D) 500 µM MNNG and (E) 300 µM etoposide. All error bars correspond to Mean±SD. Statistical significance was determined by multiple unpaired t-tests, comparing WT to *Sarm1* KO at each time point. **p < 0.01; ***p < 0.001; ****p < 0.0001.

SARM1 is the central executioner of programmed axon degeneration activated in response to a wide variety of neuronal insults^1^. We therefore investigated whether axon degeneration incited by DNA damaging agents is SARM1-dependent. Remarkably, we observed no axon degeneration in *Sarm1* KO DRG neurons treated with CPT, etoposide, or MNNG (Fig 1A-C, Supplemental Fig 1J). We tested whether DNA damage activates SARM1 by measuring cellular metabolites after treating the neurons with either etoposide or MNNG. SARM1-mediated NAD^+^ cleavage generates cyclic ADPR (cADPR), a specific biomarker of SARM1 activity^35^. Notably, cADPR levels increase approximately ∼10-fold 24 hours after addition of etoposide and ∼4-fold 8 hours after MNNG addition (Fig 1D, E). As expected, increased cADPR is not observed in *Sarm1* KO neurons. Taken together, these results demonstrate that SARM1 is activated by DNA damage and that DNA damage-induced axon degeneration is SARM1-dependent.

### DNA damage activates SARM1 via PARP1-induced NAD**^+^** depletion

Given that SARM1 is a sensor of the intracellular NMN/NAD^+^ ratio activated when NMN displaces NAD^+^ from an allosteric binding pocket^5^, we hypothesized that PARP1-mediated NAD^+^ loss is the mechanistic link between DNA damage and SARM1 activation. To test this, we first used Western blot to assess PAR, the product of PARP1 activity, and the DNA damage marker γH2AX, in neurons treated with MNNG. As expected, both γH2AX and PAR increase in MNNG-treated DRG neurons (Supplemental Fig 2A). Importantly, these increases are independent of SARM1, demonstrating that SARM1 does not suppress degeneration by blocking DNA damage or PARP1 activation, but acts downstream of these events. Next, we tested whether axon degeneration induced by MNNG is PARP1 dependent by pre-treating neurons with two structurally-distinct PARP1 inhibitors, ABT-888 (veliparib) or EB-47. Both inhibitors prevent axon degeneration in MNNG-treated neurons (Fig 2A), similar to SARM1 knockout (Fig 1B). PARP1 activation can trigger mitochondrial depolarization^36^, and we have demonstrated that mitochondrial depolarization can activate SARM1^22^, so we investigated whether this could be the mechanism of SARM1 activation. We treated neurons with MNNG in conjunction with either the PARP1 inhibitor ABT-888 or the SARM1 inhibitor NB-7^37^. Interestingly, MNNG induces a dramatic loss of mitochondrial potential within 12 hours that is blocked by either PARP1 or SARM1 inhibition (Fig 2B, C). This finding has two important implications. First, mitochondrial depolarization cannot be the mechanism of SARM1 activation since SARM1 is required for said depolarization. Second, mitochondrial depolarization after DNA damage, a hallmark of Parthanatos, is SARM1-dependent. Finally, we measured NAD^+^, NMN and cADPR, a SARM1 biomarker that is not produced by PARP1 (Fig 1D, E). MNNG leads to PARP1-dependent NAD^+^ loss (Fig 2D), and thus a PARP1-dependent increase of the NMN/NAD^+^ ratio by 6 hours post-treatment (Fig 2E). Importantly, the SARM1 biomarker cADPR also begins to rise after 6 hours of MNNG, and this increase is PARP1-dependent (Fig 2F). These results strongly support the hypothesis that DNA damage activates the PARP1 NAD^+^ hydrolase, causing an increased NMN/NAD^+^ ratio which activates SARM1 and promotes degeneration.

**Figure 2.**
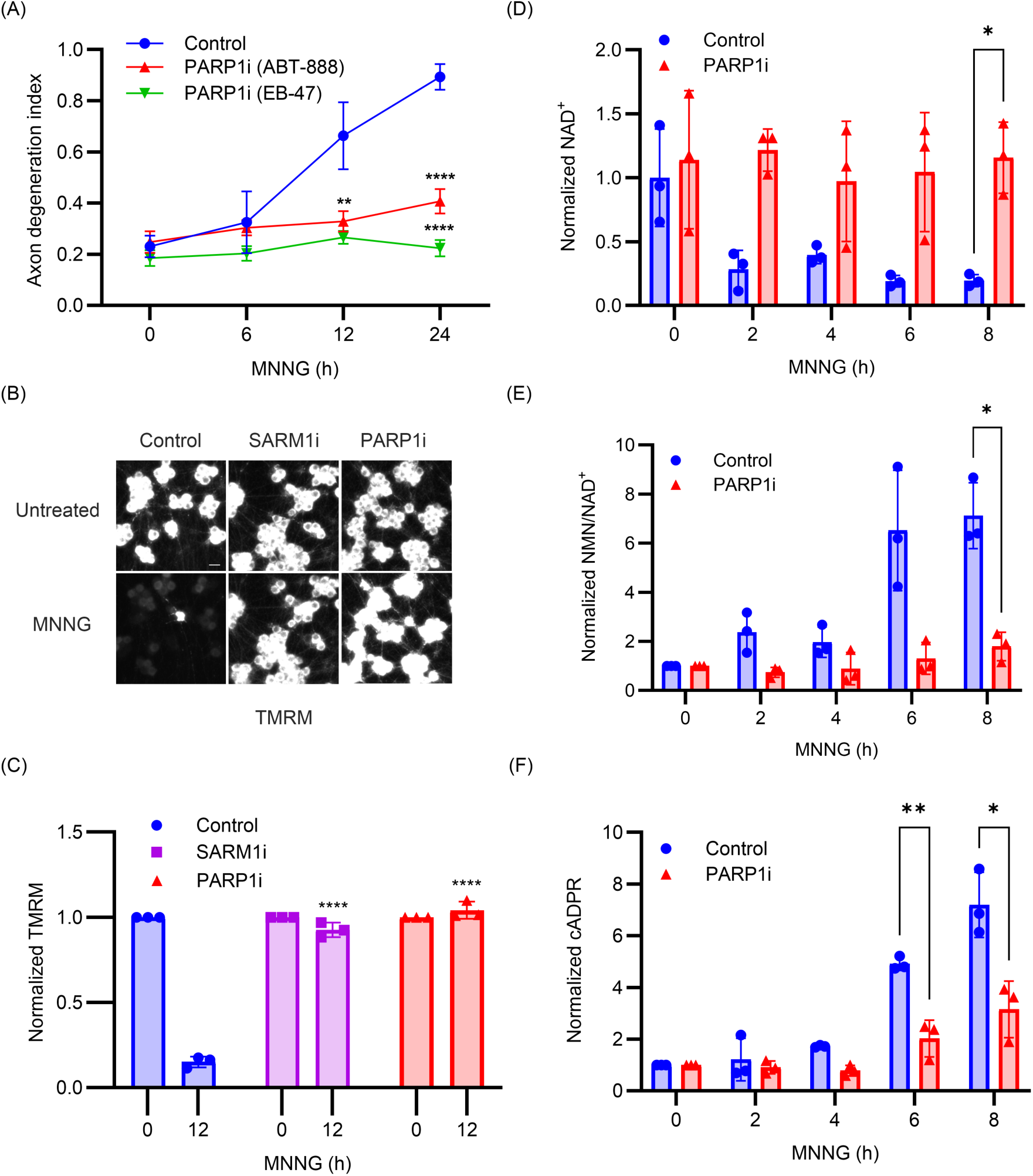
PARP1 activation is upstream of SARM1 activation. (A) DRGs preincubated with PARP1 inhibitor (ABT-888 or EB-47) were treated with MNNG (500 µM) at DIV7 and the axon degeneration index was measured. (B) DIV7 DRGs preincubated with SARM1 inhibitor (NB-7) or PARP1 inhibitor (ABT-888) were treated with MNNG (500 µM) for 12 hr and mitochondrial membrane potential assessed using TMRM dye. Scale bars, 25 µm. (C) Quantification of TMRM signal from (B). (D-F) DRGs preincubated with PARP1 inhibitor (ABT-888) were treated with MNNG (500 µM) at DIV7. Neuronal soma was collected for metabolite measurement. (D) NAD^+^, (E) NMN/NAD^+^ and (F) cADPR were quantified by LC-MS/MS. Statistical significance was determined by multiple unpaired t tests. All error bars correspond to Mean±SD. *p < 0.05; **p < 0.01; ***p < 0.001; ****p < 0.0001.

### SARM1 activation mediates AIF translocation and neuronal cell death

In Parthanatos, overactivation of PARP1 due to genomic stress leads to excessive PAR synthesis and mitochondrial dysfunction. PAR polymers interact with mitochondria and promote the release of AIF. AIF then binds MIF and translocates from mitochondria to the nucleus where they promote large-scale DNA fragmentation and chromatin condensation, ultimately leading to cell death^31^. Since SARM1 is required for mitochondrial depolarization in response to DNA damage, we examined whether SARM1 is also necessary for MNNG-stimulated neuron death and AIF translocation^38^. Approximately 50% of DRG neurons die after 12 hours exposure to MNNG (500 µM), as monitored by staining with propidium iodide (PI) assessed both microscopically (Fig 3A, B) and via flow cytometry (Supplemental Fig 3). However, SARM1 knockout neurons are fully protected from MNNG-induced cell death, as are WT neurons pre-treated with the SARM1 inhibitor NB-7 or with the PARP1 inhibitors ABT-888 or EB-47 (Fig 3A, B; Supplemental Fig 3). Hence, SARM1 is required for MNNG-mediated, PARP1-dependent cell death, i.e., SARM1 is required for Parthanatos.

**Figure 3.**
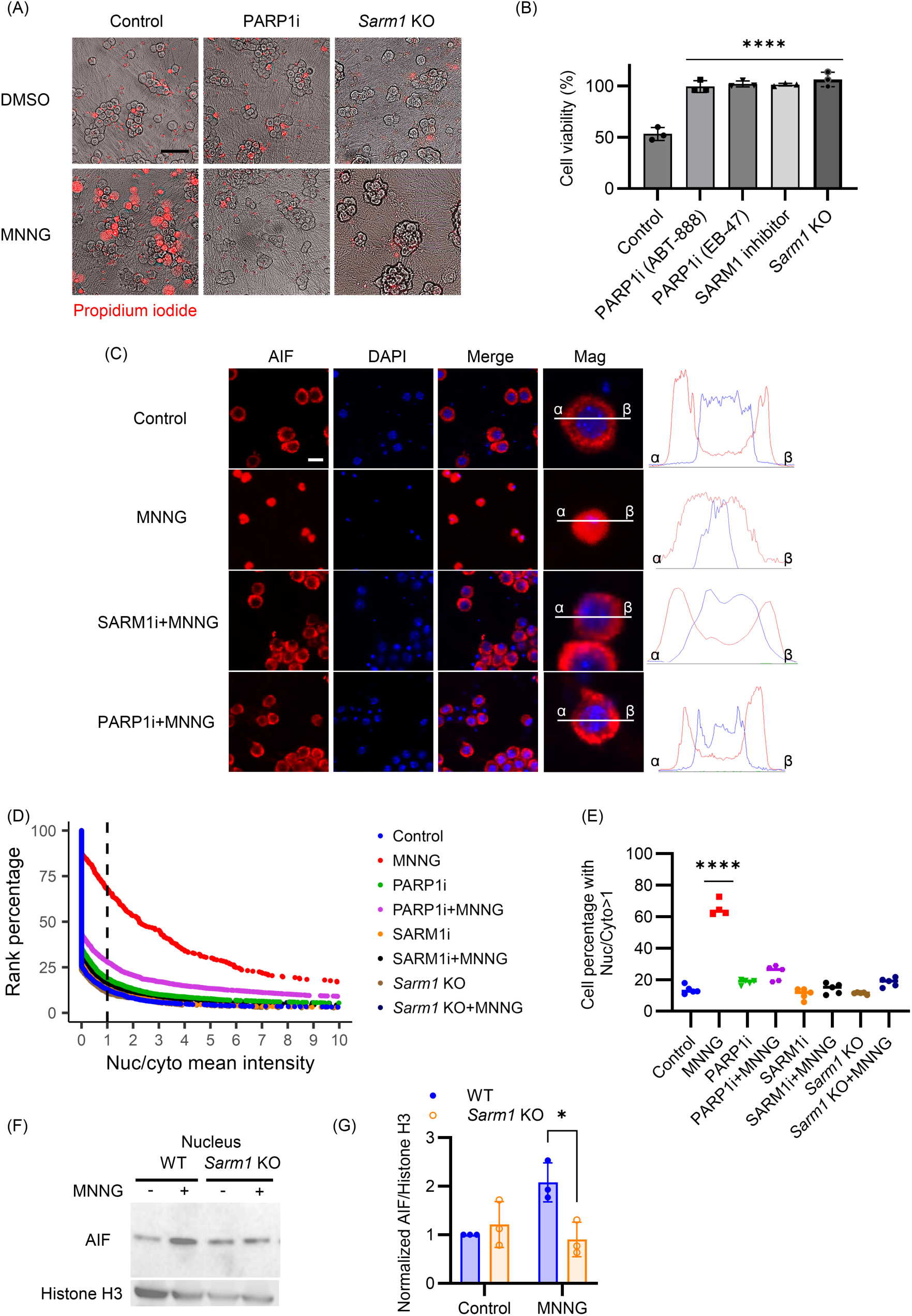
SARM1 activation induces AIF translocation and cell death. (A) WT DRG neurons, untreated or preincubated with PARP1 inhibitor (ABT-888), and *Sarm1* KO DRG neurons, were treated with MNNG (500 µM) at DIV7. Representative images show neurons after 12 hr of treatment, stained with Propidium Iodide (PI) to indicate cell death. Scale bars, 100 µm. (B) Quantification of cell viability using PI staining. WT neurons, WT neurons preincubated with SARM1 inhibitor (NB-7) or with PARP1 inhibitor (EB-47 or ABT-888), and *Sarm1* KO neurons were treated with MNNG (500 µM) at DIV7. Statistical significance was determined by one-way ANOVA, comparing all conditions to Control. (C) Representative images of immunofluorescent staining of DRG neurons with antibodies against AIF, and DAPI to label the nucleus. Neurons were preincubated with a SARM1 inhibitor or a PARP1 inhibitor, followed by treatment with 500 µM MNNG for 12 hr. AIF (red) and nuclear (blue) intensity distributions are shown for the representative magnified cell images. Scale bars, 25 µm. (D) Cumulative histogram showing the per-cell distribution of AIF intensity ratio on the x-axis, where high numbers represent more nuclear AIF compared with cytosolic. Where indicated, DRG neurons were treated with MNNG (500 µM) for 12 hr prior to quantification. (E) Quantification of percentage of cells (from D), with AIF nuclear-to-cytoplasmic intensity ratio greater than 1 (dashed line in D). Statistical significance was determined by one-way ANOVA, comparing all conditions to Control. (F) Western blot analysis shows translocation of AIF to the nucleus in WT vs. *Sarm1* KO primary cortical neurons treated with MNNG (500 µM) for 45 min. (G) Quantification of AIF intensity normalized to Histone H3 from (F). Statistical significance was determined by unpaired t tests, comparing WT and *Sarm1* KO. *p < 0.05; ****p < 0.0001.

We next explored the role of SARM1 in the signature cell biological event of Parthanatos, the translocation of AIF from mitochondria to the nucleus following genomic stress^31^. We assessed AIF translocation following MNNG treatment using immunofluorescence. Cultured DRG neurons were pretreated with SARM1 or PARP1 inhibitors, followed by MNNG treatment for 12 hours. Cells were then stained with an AIF antibody and counterstained with DAPI to visualize nuclei. In WT DRG neurons, AIF is predominantly observed in the cytosol encircling the nucleus. After MNNG treatment, much of the AIF signal overlaps with DAPI, demonstrating the translocation of AIF into the nucleus (Fig 3C). Automated quantification revealed a significant increase in the nuclear to cytosolic ratio of AIF in MNNG-treated cells (Fig 3D). As expected, AIF translocation is blocked by the PARP1 inhibitor ABT-888. Excitingly, AIF translocation is also blocked by both the SARM1 inhibitor NB-7 and in *Sarm1* KO neurons (Fig 3D, E). The SARM1-dependence of AIF translocation was also demonstrated biochemically by Western blot (Fig 3F, G). Hence, SARM1 is required for AIF nuclear translocation, the key mechanistic step of Parthanatos.

### Defective DNA repair in motor neurons harboring pathogenic *FUS* mutations promotes SARM1-dependent axon degeneration

An impaired DNA damage response leading to genomic instability in motor neurons and other cells likely contributes to the progression of ALS^39,40^. Mutations in the *Fused in Sarcoma* (*FUS*) gene cause familial ALS^41^, and FUS protein plays a critical role in DNA damage repair^42,43^. The ALS-associated *FUS*^R521H^ mutation disrupts the DNA repair complex, leading to increased DNA damage in motor neurons in both the mouse spinal cord and the human motor cortex^43,44^. To investigate the potential role of SARM1 in ALS disease progression associated with *FUS*^R521H^, we differentiated induced pluripotent stem cells (iPSCs) homozygous for the *FUS*^R521H^ mutation into motor neurons (MNs) and assessed their relative sensitivity to etoposide. We confirmed that *FUS*^R521H^ MNs are defective in DNA repair using the DNA damage marker γH2AX (Supplemental Fig 4A). Initially, the axonal area of untreated *FUS*^R521H^ MNs is identical to isogenic control (WT) MNs, but after 24 hours of etoposide, *FUS*^R521H^ MNs exhibit significantly more axon loss than similarly treated controls (Fig 4A, B). We hypothesized that the enhanced degeneration in *FUS*^R521H^ MNs is due to increased SARM1 activity. To test this, we measured NAD^+^ and its cleavage product, cADPR, in MNs treated with etoposide, assessing SARM1 activity according to the ratio of its product and substrate (cADPR/NAD^+^) as previously described^5^. *FUS*^R521H^ MNs display 2.4-fold more cADPR/NAD^+^ than isogenic controls after 48 hours treatment with etoposide (p = 0.03, n=3). Next, we assessed the requirement for SARM1 in mediating degeneration in response to DNA damage by pretreating *FUS*^R521H^ MNs with the SARM1 inhibitor NB-7. Metabolite analysis demonstrates that NB-7 prevents the increase of cADPR/NAD^+^ in *FUS*^R521H^ MNs after etoposide treatment (Fig 4C), paralleling a significant protection from axonal degeneration (Fig 4A, B). Moreover, SARM1 inhibition results in the axonal area of etoposide-treated *FUS*^R521H^ MNs being indistinguishable from the similarly treated controls, demonstrating that the enhanced etoposide sensitivity of *FUS*^R521H^ MNs is fully SARM1-dependent. In conclusion, SARM1 contributes to DNA damage-induced motor axon pathology in the context of *FUS* mutation, suggesting that SARM1 inhibition could mitigate the consequences of DNA damage associated with ALS.

**Figure 4.**
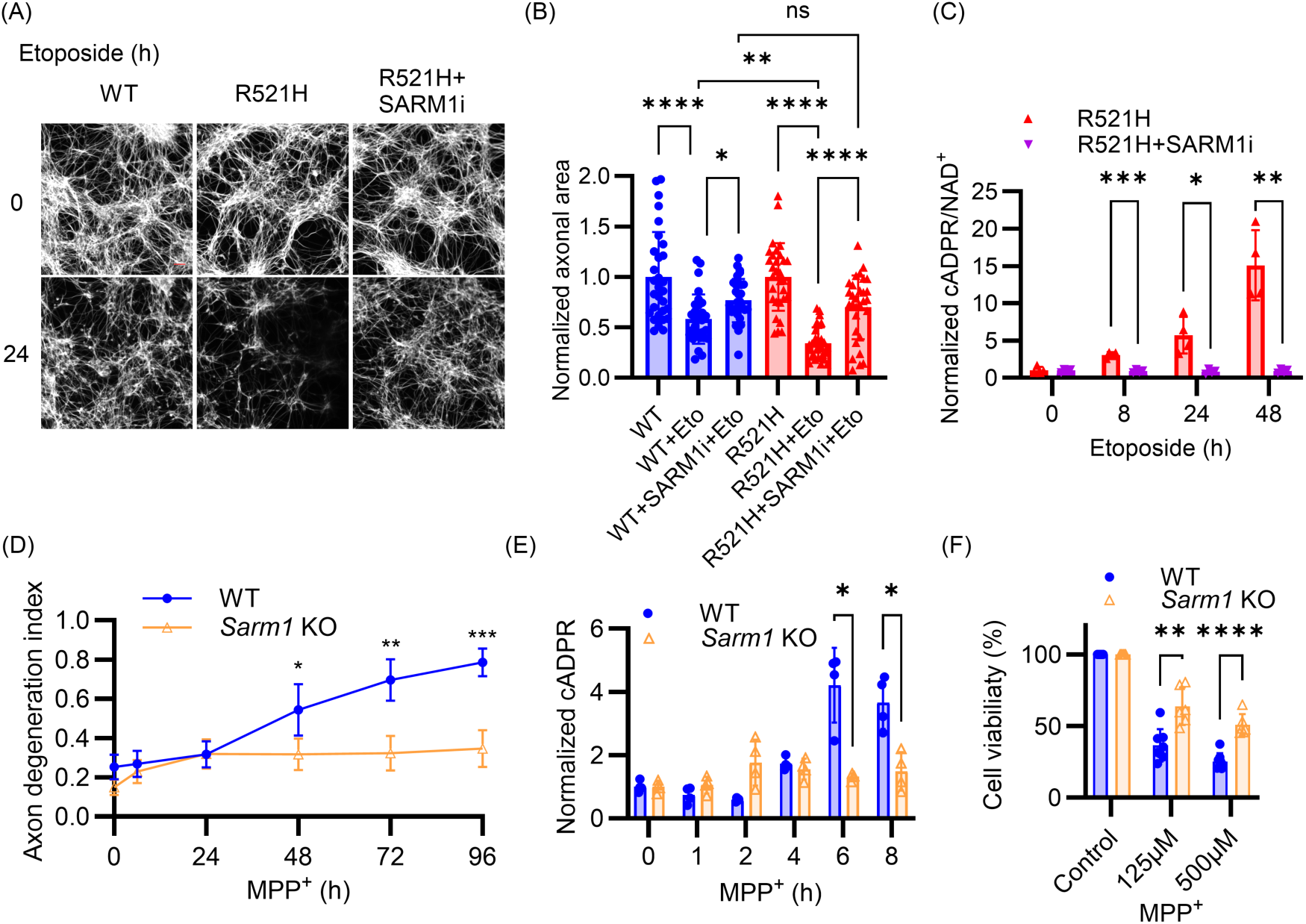
SARM1 inhibition prevents neuronal pathology associated with *FUS* mutation and MPP^+^ toxicity. (A) Representative images of axons from WT and *FUS*^R521H^ iPSC-derived motor neurons treated with 10 µM etoposide, stained with Tubulin Tracker™ Deep Red at 0 and 24 hr. Scale bars: 50 µm. (B) Quantification of axonal area in *FUS*^R521H^ and isogenic control (WT) iPSC-derived MNs 24 hr after treatment with 10 µM etoposide (Eto) and SARM1 inhibitor (NB-7). Statistical significance was determined by one-way ANOVA. (C) *FUS^R521H^* iPSC-derived motor neurons preincubated with SARM1 inhibitor (NB-7) were treated with etoposide (10 µM), and metabolites collected. cADPR/NAD^+^ was quantified by LC-MS/MS. Statistical significance was determined by multiple unpaired t tests comparing inhibitor-treated to untreated at each time point. (D) WT and *Sarm1* KO DRG neurons were treated with 1 mM MPP^+^ at DIV7 and axon degeneration index measured at indicated times post-treatment. (E) WT and *Sarm1* KO DRG neurons were treated with 100 µM MPP^+^ at DIV7 and cADPR quantified by LC-MS/MS at indicated times after treatment. (F) WT and *Sarm1* KO DRG neurons were treated with indicated doses of MPP^+^ for 48 hr at DIV 7 and MTT assays were performed to quantify cell viability. Statistical significance in (D-F) was determined by multiple unpaired t tests, comparing *Sarm1* KO to WT at each time point or each dose. All data and error bars correspond to Mean±SD. *p < 0.05; **p < 0.01; ***p < 0.001; ****p < 0.0001.

### SARM1 promotes MPP^+^ neurotoxicity

Parthanatos is a key driver of neurodegeneration in Parkinson’s disease, and inhibition of Parthanatos protects against 1-methyl-4-phenyl-1,2,3,6-tetrahydropyridine (MPTP) neurotoxicity *in vivo*^32^, a commonly used model of parkinsonism. Given the crucial role of PARP1 activation in MPTP neurotoxicity and MPTP-induced parkinsonism^45,46^, we hypothesized that SARM1 activation contributes to MPTP-induced neurodegeneration. MPTP requires metabolic conversion to MPP^+^ by glial cells or astrocytes to exhibit its toxicity^47^. We therefore used MPP^+^ iodide to treat DRG neurons *in vitro*. Excitingly, MPP^+^-induced axon degeneration is fully SARM1-dependent (Fig 4D). Furthermore, MPP^+^ treatment leads to a SARM1-dependent decrease in NAD^+^ and increase in cADPR, and a decrease in ATP that is partially SARM1-dependent (Fig 4E, and Supplemental Fig 4B, C). Importantly, MPP^+^-treated *Sarm1* KO neurons display significantly less cell death than WT neurons (Fig 4F). Thus, SARM1 inhibition protects against axon degeneration and partially prevents neuron death induced by MPP^+^.

### SARM1 promotes NMDA excitotoxicity

Parthanatos-mediated neuron death is central to glutamate excitotoxicity, a pathological process implicated in many neurodegenerative diseases and ischemic stroke^48–50^. Excessive NMDA receptor (NMDAR) activation leads to an influx of Ca²⁺, excessive ROS generation, and subsequent DNA damage, ultimately triggering cell death^49,51^. Since SARM1 is required for Parthanatos in neurons, we investigated whether SARM1 inhibition protects against excitotoxicity. We exposed primary cortical neurons to NMDA for 24 hours, with or without SARM1 inhibitor pretreatment, and assessed neuronal viability using NeuO, a dye that selectively stains live neurons^52,53^. The inhibition of SARM1 significantly protects neurons from NMDA-induced death at a level comparable to the NMDAR antagonist MK801^54,55^ (Fig. 5A). Excitotoxicity is also associated with mitochondrial dysfunction, particularly the loss of mitochondrial membrane potential^56,57^, and SARM1 inhibition prevents NMDA-induced mitochondrial depolarization (Fig. 5B). Beyond cell survival, maintaining functional neuronal connectivity is crucial for preserving neural circuit integrity. We therefore assessed neurite degeneration following NMDA exposure as an indicator of functional deterioration. SARM1 inhibition, like MK801, fully preserves neurite integrity despite excitotoxic stress (Fig. 5C).

**Figure 5.**
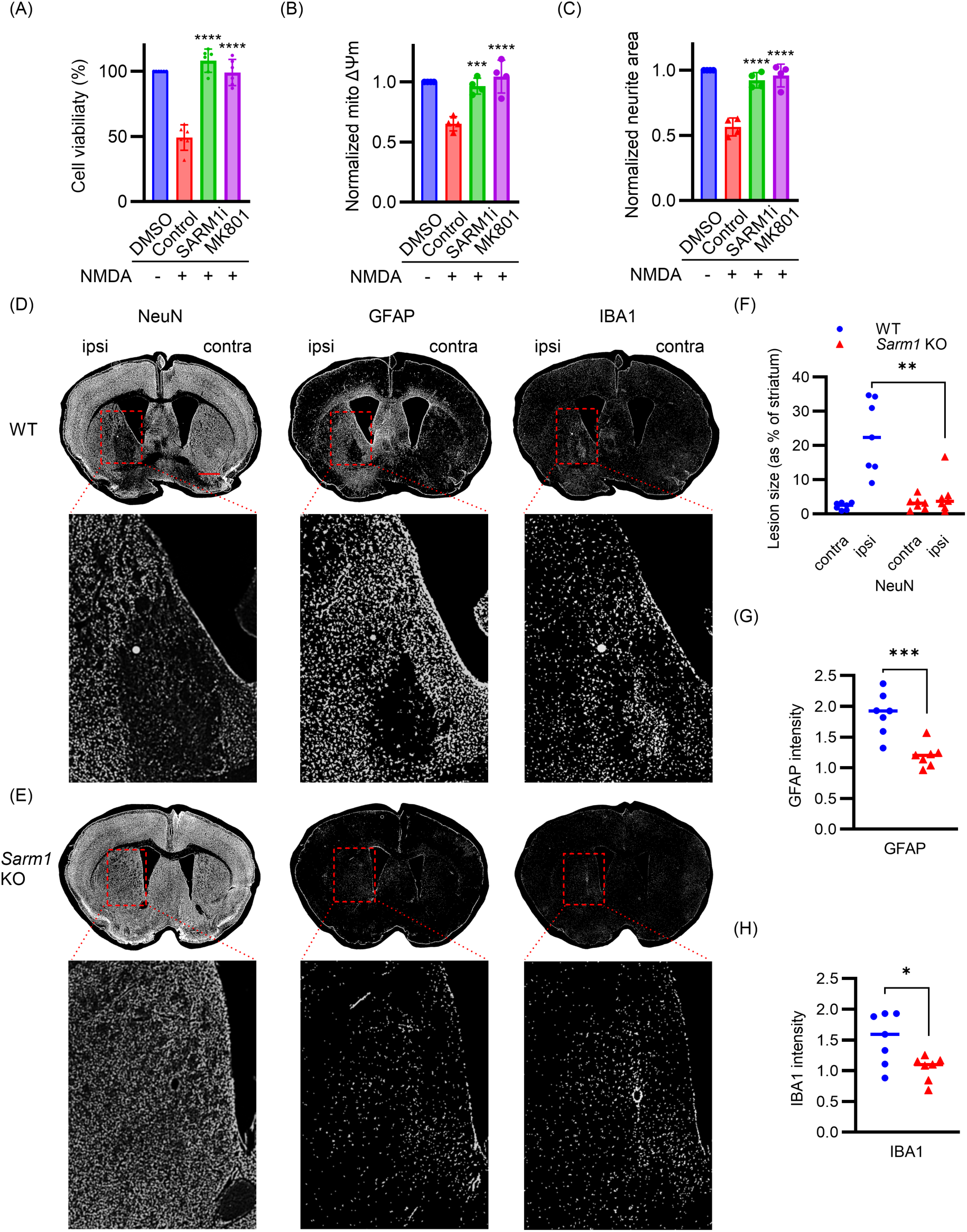
SARM1 inhibition protects neurons from excitotoxicity *in vitro* and *in vivo*. (A) Quantification of cell viability by averaging NeuO intensity. Primary cortical neurons preincubated with DMSO, the SARM1 inhibitor NB-7 or the NMDAR antagonist MK801 were treated with NMDA (500 µM) at DIV14. Cell viability was quantified after 24 hours of NMDA stimulus. (B) Quantification of mitochondrial membrane potential by measuring average TMRM intensity in primary cortical neurons after 24 hours of NMDA stimulus. (C) Quantification of neurite area in primary cortical neurons after 24 hours of NMDA stimulus. Statistical significance in (A-C) was determined by one-way ANOVA, comparing NMDA control with SARM1 inhibitor or MK801 preincubation. (D-E) WT and *Sarm1* KO Mice were injected intrastriatally with NMDA and tissues collected after 48 hr. (D) and (E) show representative images of coronal brain sections from WT (D) and *Sarm1* KO (E) mice immunostained for NeuN (neurons), GFAP (astrocytes), and IBA1 (microglia). Scale bars, 1.5 mm. (F) Quantification of lesion area in the striatum, calculated as the percentage of NeuN-depleted area relative to the total striatal area per hemisphere. Lesion and total areas were determined using a threshold-based grid quantification of NeuN intensity. (G-H) Fold increase of (G) GFAP and (H) IBA1 intensity between the ipsilateral and contralateral striatum for each mouse. Statistical significance in (F-H) was determined by multiple unpaired t tests. *p < 0.05; **p < 0.01; ***p < 0.001; ****p < 0.0001.

To assess the role of SARM1 in excitotoxicity *in vivo*, we injected NMDA into the striatum of WT and *Sarm1* KO mice. Brain sections were analyzed 48 hours post-injection to evaluate neuronal loss, gliosis and neuroinflammation by staining for the neuronal marker NeuN, the activated astrocyte marker GFAP and the microglial marker IBA1. In WT mice, NMDA injection resulted in large lesions with substantial NeuN loss (Fig. 5D). In contrast, neurons were significantly retained in *Sarm1* KO mice (Fig. 5E). GFAP expression was also significantly higher in the ipsilateral striatum of WT brains compared to the contralateral, except in the core of the lesion where astrocyte loss was apparent (Fig. 5D). IBA1 staining was elevated within the lesion, indicating robust microglial activation at the site of degeneration. By comparison, *Sarm1* KO brains showed similar GFAP and IBA1 expression on their ipsilateral and contralateral sides (Fig. 5E). Quantitative analysis confirmed significant neuronal loss, astrocyte activation, and microglial recruitment in the ipsilateral striatum of WT mice, which were markedly attenuated in *Sarm1* KO animals (Fig. 5F–H). Hence, SARM1 promotes NMDA-induced excitotoxic injury *in vivo*, highlighting the potential of SARM1 inhibition as a therapeutic strategy for excitotoxic neurodegenerative diseases.

## DISCUSSION

The role of SARM1 as the central executioner of axon degeneration is well understood^3,5^, but the mechanisms by which it can be activated to induce neuronal cell death^1,58^ had not been defined. Here, we demonstrate that SARM1 is a heretofore unappreciated essential component of neuronal Parthanatos activated when insults such as DNA damaging agents stimulate PARP1 NAD^+^ hydrolysis, raising the NMN/NAD^+^ ratio. The clinical implications of this finding are significant, providing a mechanistic rationale for testing SARM1 inhibitors as treatments for Parthanatos-associated diseases such as Parkinson’s and stroke.

SARM1 triggers neurodegeneration by depleting cellular NAD^+^, while Parthanatos is understood to drive cell death through the cytoplasmic accumulation of PAR. Therefore, the two pathways have been viewed as distinct. However, Parthanatos and sarmoptosis have two crucial commonalities—both are independent of other well-defined cell death pathways i.e. unaffected by inhibitors of apoptosis or necroptosis^59,60^, and both pathways involve significant NAD^+^ depletion via their central NAD^+^-hydrolases, PARP1 and SARM1. Reducing intracellular NAD^+^ is sufficient to activate SARM1^5^, and thus the dramatic NAD^+^ loss following PARP1 overactivation^61^ suggests an obvious mechanistic bridge to SARM1. Not surprisingly, investigators originally proposed that PARP1 effectuated cell death by precipitating metabolic collapse^62^, and more recently, links between SARM1 and PARP1 have been investigated^63^. Nevertheless, the field has largely turned away from that NAD^+^ depletion hypothesis for Parthanatos. Our study forces reevaluation of that conclusion and reinterpretation of prior findings.

According to the current standard model of Parthanatos, excessive DNA damage hyperactivates PARP1, which consumes NAD^+^ to polymerize PAR. PAR is transported to the cytoplasm where it binds to Hexokinase1 and AIF, interactions that respectively inhibit glycolysis and activate nuclease activities that result in DNA fragmentation. These changes are occasioned by sharply decreased cellular NAD^+^ and ATP^64^. More than forty years ago, it was found that PARP1 inhibition or supplementation with NAD^+^ precursors could preserve NAD^+^ and ATP and maintain cellular metabolism after DNA damage^65,66^. Understandably, the NAD^+^ loss was attributed directly to PARP hydrolase activity, an assumption supported by later work demonstrating that NAD^+^ supplementation prevents mitochondrial permeability transition and AIF nuclear translocation^36^. Conversely, introduction of an NAD^+^ hydrolase is sufficient to trigger AIF translocation and cell death in wild type and *PARP1^-/-^*neurons^67^. However, two seminal papers later demonstrated that blocking NAD^+^ synthesis with FK866, a highly specific inhibitor of nicotinamide phosphoribosyltrasferase (NAMPT), could significantly deplete NAD^+^ without affecting ATP levels, glycolysis or mitochondrial function^28,68^, suggesting that mechanisms other than NAD^+^ loss are more important for Parthanatos. Thereafter, the field refocused on PAR and its interactions with extranuclear factors. At the time, no one could have accounted for SARM1 and its unique activation mechanism.

We have now demonstrated, via both genetic knockout and direct chemical inhibition, that the NAD^+^ hydrolase activity of SARM1 is necessary to execute Parthanatos in neurons. Under normal cellular conditions, significant PARP1-mediated NAD^+^ depletion activates SARM1, a voracious hydrolase that contributes to NAD^+^ depletion when the NMN/NAD^+^ ratio rises. Therefore, prior experiments examining PARP1-dependent cell death in neurons should be reinterpreted in light of SARM1. Notably, we recently demonstrated that SARM1 activation is regulated by the competitive binding of NAD^+^ and its precursor NMN to an allosteric site on the enzyme^5^. NAD^+^ is synthesized from nicotinamide by the sequential action of NAMPT, which generates NMN, and NMNAT enzymes, which convert NMN to NAD^+^. When NMNAT activity is reduced, NMN accumulates and outcompetes NAD^+^ to activate SARM1. However, inhibition of NAMPT reduces the levels of both NMN and NAD^+^ and so does not activate SARM1^5^. Hence, the prior efforts to test the role of NAD^+^ depletion in Parthanatos by treating with the NAMPT inhibitor FK866 were confounded by the concomitant loss of NMN.

PARP1 and SARM1 are both necessary for Parthanatos, and abundant evidence indicates that excessive PAR generation contributes to cell death^24,31^. We propose that PAR toxicity and NAD^+^ loss collaborate to kill neurons in Parthanatos via the following mechanism: DNA damage activates PARP1, which both depletes NAD^+^ and generates PAR. This drop in NAD^+^ activates SARM1, which further depletes NAD^+^ and induces mitochondrial depolarization, a well-described function of active SARM1^10,19^. Mitochondrial damage makes AIF accessible to PAR, and PAR-AIF binding then facilitates AIF/MIF nuclear translocation leading to massive DNA fragmentation. This proposed mechanism brings together diverse experimental evidence regarding DNA damage-induced neuronal death. However, we expect that the relative contributions of canonical Parthanatos components and SARM1 varies by cell type and pathological insult.

The SARM1 hydrolase is a conspicuously *druggable* therapeutic target with inhibitors already in clinical trials^69–71^. While the standard model of Parthanatos has potential therapeutic points of entry, they are not without complications. Inhibiting PARP1 potently blocks Parthanatos, but PARP1 is required for DNA damage repair and blocking only “excess” PARP1 activity is challenging. Strategies to block the MIF nuclease also hold promise, but DNA fragmentation is the *last* step in Parthanatos, downstream of NAD^+^ depletion and mitochondrial depolarization, both of which seriously undermine cellular viability^32^. In contrast, SARM1 functions upstream of mitochondrial depolarization and AIF nuclear translocation. Thus, targeting SARM1 to halt Parthanatos would boost cellular metabolite homeostasis, preserve mitochondrial integrity, and prevent MIF-mediated DNA damage. Notably, SARM1 loss also protects from neurodegeneration by preserving mitochondria in other scenarios such as congenital CMT2A^14^. As such, the pivotal role of SARM1 in executing neuronal Parthanatos dramatically expands the potential repertoire of SARM1-targetting therapies to include most common neurodegenerative diseases, especially those associated with neuronal excitotoxicity—Parkinson’s, Alzheimer’s, ALS, and Huntington’s, as well as stroke^24,31,48,59,72^. Indeed, this potential is highlighted by the rescue of neuropathology following striatal injection of NMDA into *Sarm1* KO animals shown above. A clear path for blocking Parthanatos provides a new hope for confronting these devastating ailments.

## DECLARATIONS

## Author contributions

TW, YS, AJB, AD, JM conceived the overall study and study design. TW and LY performed the experiments. TW and WB analyzed the data and prepared figures. TW prepared the manuscript. YS, AJB, AD, JM oversaw the analysis and edited the manuscript.

## Acknowledgements

We thank Milbrandt and DiAntonio lab members for technical support and insightful comments during this study, especially Cassidy Menendez, Waleed Minzal and Ana Morales Benitez. Thanks to Amber Neilson, Xiaoxia Cui, Jason Waligorski, Colin Kremitzki for providing iPSC resources at the McDonnell Genome Institute. Thanks also to Susie Grathwohl at the Hope Center Animal Surgery Core.

## MATERIALS AND METHODS

### Cell culture

HEK 293T cells were cultured and passed in Dulbecco’s Modified Eagle Medium (DMEM) (Thermofisher Scientific) supplemented with 10% FBS, glutamine and penicillin/streptomycin. Cells were cultured in a 37 °C incubator with 5% CO_2_. Cells were passed every 3-4 days by using 0.05% Trypsin containing 0.02% EDTA (Gibco) and transferring 1:10 volume to a new T75 tissue culture flask.

### Mice

All experiments were conducted in accordance with the protocols approved by the Institutional Animal Care and Use Committee (IACUC) of Washington University in St. Louis and adhered to the National Institutes of Health (NIH) Guidelines for the Care and Use of Laboratory Animals. Mice were maintained on a 12-hour light-dark cycle, with no more than five animals per cage, and had ad libitum access to food and water. Both male and female mice were included in all experiments.

### DRG neuron culture and metabolite extraction

Mouse DRG cultures were prepared following established protocols^73^. C57BL/6 or *Sarm1* KO pregnant mice were sacrificed at embryonic day 13.5 (E13.5) and approximately 50 DRGs harvested per embryo. The ganglia were dissociated in a solution containing 0.05% Trypsin and 0.02% EDTA (Gibco) at 37 °C for 20 minutes. Following digestion, cells were washed three times with DRG culture media to remove residual trypsin. Dissociated neurons were plated as spot cultures in 96-well tissue culture plates (Corning) or 24-well tissue culture plates (Corning) pre-coated with 0.1 mg/ml poly-D-lysine and 0.1 mg/ml laminin. Cells were maintained in Neurobasal medium (Gibco) supplemented with 2% B27 (Invitrogen), 100 ng/ml 2.5S NGF (Harlan Bioproducts), and 1 μM 5-fluoro-2′-deoxyuridine/1 μM uridine. Media was refreshed every 2–3 days. Lentiviral transduction was performed on day in vitro 3 (DIV3). For metabolite analysis, culture medium was aspirated from neurons grown in 24-well tissue culture plates, and 160 μL of ice-cold 50% methanol (MeOH) was added to each well to extract cellular metabolites. Plates were incubated at 4 °C for 20 minutes, after which the extract from each well was collected and mixed with 40 μL of chloroform (Sigma-Aldrich). The mixture was centrifuged at 15,000 rpm for 10 minutes, and 140 μL of the upper aqueous phase was carefully collected for further processing. Samples were then lyophilized using a SpeedVac™ Medium Capacity Concentrator (Thermo Fisher Scientific) and stored or reconstituted for downstream metabolomic assays. For experiments involving PARP1 inhibitor pretreatment, the central region of spot cultures surrounding neuronal somas was harvested using a sterile razor blade to enrich for soma-derived metabolites. Unless otherwise specified, all other metabolite measurements were performed using whole cells.

### Primary cortical neuron culture

Cortical tissue was isolated from embryonic day 14.5 (E14.5) C57BL/6 embryos and enzymatically dissociated using 0.05% trypsin at 37°C for 18 minutes. The dissociated neurons were subsequently washed in culture medium [Neurobasal (21103; Gibco) supplemented with 2% B27 (Invitrogen), 1 μM 5-fluoro-2′-deoxyuridine, and 1 μM uridine]. Cells were then seeded at a density of 6 × 10^4^ per well in 96-well plates pre-coated with 0.1 mg/ml poly-D-lysine. Culture medium was refreshed every four days with 50% replaced at each interval. Drug treatment

DRGs neurons were pretreated for 1 hour with 25 µM NB-7 for SARM1 inhibition. PARP1 inhibitors, including ABT-888 (Cayman Chemical Company) and EB-47 (MedChem Express), were applied at a concentration of 100 µM overnight. To induce axon degeneration, DRG neurons were treated with either 10 µM camptothecin (CPT, Sigma-Aldrich), 300 µM etoposide (Selleck Chemicals) or 500 µM MNNG (Selleck Chemicals).

### Cell sorting

After 12 hours of MNNG treatment, DRG neurons were stained with Hoechst and Propidium Iodide for 15 minutes. Cells were then analyzed using a CytoFLEX Flow Cytometer (Beckman), with approximately 2 × 10⁶ events recorded. Cell death was quantified using the V450-A and B690-A channels, with double-positive cells classified as dead.

### iPSC differentiation protocol

Human iPSCs were differentiated into motor neurons following established protocols with slight modifications^74^. Briefly, iPSCs were maintained in a chemically defined neural medium containing DMEM/F12 and Neurobasal at a 1:1 ratio, supplemented with N2, B27, ascorbic acid, GlutaMAX, and penicillin/streptomycin. Neural induction was achieved using CHIR99021, DMH1, and SB431542, followed by patterning with retinoic acid and purmorphamine to generate OLIG2^+^ motor neuron progenitors (MNPs). MNPs were expanded and subsequently differentiated into motor neurons with additional treatment of valproic acid. Maturation into CHAT^+^ motor neurons was facilitated by Compound E and neurotrophic factors, including IGF-1, BDNF, and CNTF. Both WT and *FUS^R521H^* homozygous mutant iPSCs were obtained from JAX iPS Cells (Jackson Laboratory). To assess axon degeneration and metabolite changes, cells were treated with 10 µM etoposide at the mature motor neuron stage.

### AIF translocation and Immunofluorescence

Following 12 hours of MNNG treatment, neurons were washed with DPBS (Thermo Fisher Scientific), fixed with 4% paraformaldehyde for 15 minutes, and stored in DPBS for staining. Cells were permeabilized with 0.3% Triton X-100 and incubated overnight with a 1:400 dilution of AIF antibody (D39D2, Cell Signaling). Secondary antibody using Goat anti-rabbit Cy3 conjugated (1:1000; 111-165-144, Jackson ImmunoResearch) for 1 hour at room temperature. Staining was performed alongside DAPI counterstaining for nuclear visualization. Cellular images were acquired using the IN Cell Analyzer 6500, and AIF translocation was quantified with IN Carta Image Analysis Software. The cytoplasm was estimated with a 3 μm ’collar’ in InCarta, and the background-subtracted intensity of AIF was measured automatically in the cytoplasmic and nuclear masks. The small collar was key to limit the amount of background and stay focused on the small cell body area outside of the nuclei. Data for each individual cells was analyzed with Tibco Spotfire Analyst, where cells were filtered or gated to exclude debris and dead cells (nuclei area between 72 and 250, nuclei intensity between 555 and 1800), as well as to only include cells that were AIF positive.

### Antibodies and western blot

Phospho-Histone H2A.X (Ser139) (20E3) (Cell Signaling) and Poly/Mono-ADP Ribose (D9P7Z) (Cell Signaling), GAPDH (D16H11) (Cell Signaling), β3-Tubulin (D71G9) (Cell Signaling), and β-actin (Sigma-Aldrich) antibodies were used to access DNA damage in cells. All antibodies were diluted with 1:1000 ratio for western blot.

### Excitotoxicity assay

Primary cortical neurons cultured in 96-well plates at DIV14 were used for NMDA stimulation. Culture medium was carefully removed, and cells were incubated with control salt solution (CSS) [120 mM NaCl, 5.4 mM KCl, 1.8 mM CaCl₂, 25 mM Tris-HCl (pH 7.4), and 15 mM D-glucose] for 5 minutes, following established protocols^49,75^. To induce excitotoxicity, 500 µM NMDA and 10 µM glycine were added to CSS. After a 5-minute incubation, the CSS was replaced with fresh culture medium and maintained for 24 hours. Care was taken to minimize drying of the cells during medium exchange. To assess the effects of pharmacological treatments, 25 µM SARM1 inhibitor NB-7 or 10 µM NMDA receptor antagonist MK801 was added to the culture medium 1 hour prior to NMDA stimulation, and the same concentrations were maintained in the replacing medium. After 24 hours of NMDA exposure, cells were incubated with NeuO (STEMCELL Technologies), TMRM dye, Tubulin Tracker™ Deep Red (Thermo Fisher Scientific), and Hoechst for 30 minutes before imaging. Cellular images were acquired using the IN Cell Analyzer 6500, and image analysis was performed with IN Carta Image Analysis Software. Hoechst staining was used to generate nuclear masks, while NeuO, TMRM, and Tubulin Tracker™ Deep Red staining were used to create respective masks for neuronal cell bodies, mitochondrial membrane potential, and neurite structures. The software algorithm calculated the average NeuO intensity (indicating cell viability), average TMRM intensity (reflecting mitochondrial membrane potential), and the average tubulin-stained area per cell (representing neurite area).

### NMDA intrastriatal injection

Mice aged approximately 8 months were anesthetized by intraperitoneal injection of pentobarbital sodium (85 mg/kg; Sagent Pharmaceuticals). Following a midline scalp incision, a burr hole was drilled at stereotaxic coordinates: 0.5 mm rostral, 1.7 mm lateral, and 3.5 mm ventral from bregma. A total of 20 nmol NMDA was delivered into the striatum (based on a ∼20 g body weight) using a digital stereotaxic frame and a Nanomite Injector Syringe Pump. The injection was followed by a 10-minute dwell time, with an additional 3-minute pause before needle withdrawal to facilitate proper diffusion. Post-injection, mice were placed in clean cages within a 37 °C incubator and supplied with supplemental oxygen to prevent hypoxia. Animals typically recovered within 90–120 minutes. After 48 hours post-injection, mice were anesthetized with isoflurane and perfused with ice-cold PBS followed by 4% paraformaldehyde (PFA) in PBS. Brains were post-fixed overnight in 4% PFA, cryoprotected in 30% sucrose/PBS, then embedded in O.C.T (Fisher HealthCare) and sectioned at 40 μm thickness using a Cryostat. Coronal brain sections were incubated with the following primary antibodies for 1 hour at room temperature: anti-NeuN (1:200; ABN91, Millipore), anti-IBA1 (1:200; 019-19741, FUJIFILM Wako). Secondary antibodies included anti-chicken Cy3 (703-165-155, Jackson ImmunoResearch) and anti-rabbit Alexa Fluor 647 (111-605-003, Jackson ImmunoResearch). GFAP was detected using a directly conjugated Alexa Fluor 488 monoclonal antibody (1:250; GA5, Thermo Fisher Scientific), incubated for 1 hour at room temperature. To quantify lesion size following NMDA injection, NeuN-stained coronal brain sections were analyzed. The striatum was manually cropped from each image based on anatomical boundaries defined by DAPI counterstaining. All cropped images were standardized to the same dimensions to ensure consistent analysis. Image analysis was performed using ImageJ software (NIH) with a custom macro called “Grid quantification”. This macro divides each image into a 50 × 50 grid (totaling 2500 regions of interest, ROIs) and calculates the mean gray value for each grid square. The data for each grid are stored in the ROI Manager and exported for further analysis. To calculate the lesion area, background grids outside the striatal region were excluded, and a gray value threshold was applied to distinguish lesioned regions with lower NeuN intensity from intact tissue. The percentage of lesioned area was calculated as the number of grids below the NeuN intensity threshold divided by the total number of valid striatal grids. This value reflects the proportion of the striatum affected by neuronal loss. The “Grid quantification” macro was originally developed and was available in this paper. Quantifications of GFAP and IBA1 were the ratios of intensity between the ipsilateral and contralateral striatum for each mouse.

### Quantification of axon degeneration

After drug treatment to induce axon degeneration, bright-field images of the distal axons were then captured using IN Cell Analyzer 6500. Axon degeneration was quantified using ImageJ (NIH), with the degeneration index determined as the ratio of fragmented axon areas, following previously established methods with modification^73^. For each condition, the degeneration index was averaged across 6 fields per well.

### Metabolite measurement using LC-MS/MS

Lyophilized samples were reconstituted in 15 μl of 5 mM ammonium formate and centrifuged at 12,000 × g for 10 minutes. The cleared supernatants were transferred to sample vials. Calibration curves were generated using serial dilutions of metabolite standards in 5 mM ammonium formate. High-performance liquid chromatography-mass spectrometry (HPLC-MS) analysis was conducted using an Agilent 1290 Infinity II liquid chromatography system (Agilent Technologies, Santa Clara, CA) and Atlantis T3 column (2.1 × 150 mm, 3 μm) with a VanGuard guard cartridge (2.1 × 5 mm, 3 μm) (Waters, Milford, MA). The system was coupled to an Agilent 6470 Triple Quadrupole mass spectrometer (Agilent Technologies, Santa Clara, CA). Chromatographic separation was performed at a flow rate of 0.15 ml/min using a mobile phase composed of 5 mM ammonium formate in water (solvent A) and 100% methanol (solvent B). The column was equilibrated with 0% B, maintained for 2 minutes post-injection, followed by a linear gradient to 20% B over 4 minutes. The gradient was further ramped to 50% B over 2 minutes and held for an additional 2 minutes before reverting to 0% B over 5 minutes, with re-equilibration at 0% B for 9 minutes. The total run time per sample was 24 minutes, with an injection volume of 10 μl. The mass spectrometer, equipped with an electrospray ionization (ESI) source, was operated in positive ion multiple reaction monitoring (MRM) mode for metabolite detection. The [M+H]+ transitions were optimized for each metabolite as follows: NAD+ (m/z 664 → 428), NMN (m/z 335 → 123), cADPR (m/z 542 → 428), ATP (m/z 508 → 136). The fragmentation, collision energy (CE), and cell accelerator voltage were optimized for each transition. Data acquisition and quantification were performed using MassHunter Workstation software version B.08.00 for the 6400 Series Triple Quadrupole (Agilent Technologies, Santa Clara, CA).

### Quantification and statistical analysis

Statistical analyses were conducted using one-way analysis of variance (ANOVA) for comparisons among multiple groups, while two-tailed t-tests were used for pairwise comparisons to determine statistical significance. Significance was assessed based on p-values, with thresholds defined as follows: *p < 0.05; **p < 0.01; ***p < 0.001; ****p < 0.0001. All error bars indicate standard deviation (SD). Axon degeneration indexes were quantified from bright-field images of distal axons using an ImageJ macro as previously described with modification^73^. Data visualization, including line graphs, bar graphs and dot graphs, was performed using GraphPad Prism 7 and R. Detailed statistical information for each experiment is provided in the corresponding figure legends.

---

**Supplemental Fig. 1:**
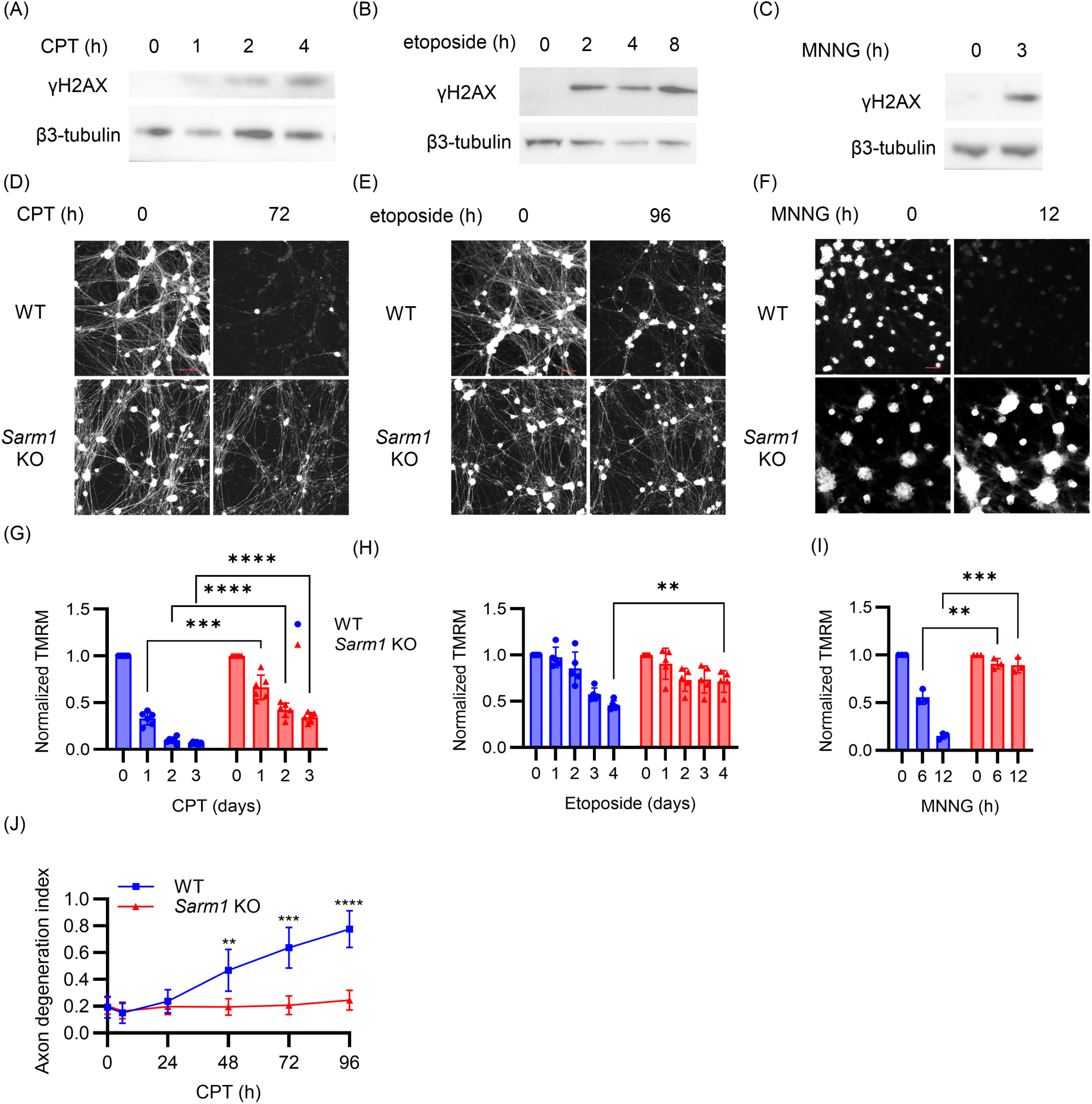
(A-C) Western blot analysis of the DNA damage marker γH2AX in WT and *Sarm1* KO DRG neurons treated with (A) CPT (10 µM), (B) etoposide (300 µM) and (C) MNNG (500 µM) at indicated time points. (D-F) WT and *Sarm1* KO DRG neurons were treated with (D) CPT (10 µM), (E) etoposide (300 µM) and (F) MNNG (500 µM) at DIV7 and stained with TMRM dye to assess mitochondrial membrane potential. Scale bars, 100 µm. (G-I) Quantification of TMRM signal from images above of WT and *Sarm1* KO DRG neurons after treatment with (G) CPT (10 µM), (H) etoposide (300 µM) and (I) MNNG (500 µM) at DIV7. (J) Axon degeneration index after treatment with 10 µM CPT. Statistical significance was determined by multiple unpaired t tests, comparing WT to *Sarm1* KO at each time point. **p < 0.01; ***p < 0.001; ****p < 0.0001.

**Supplemental Fig. 2:**
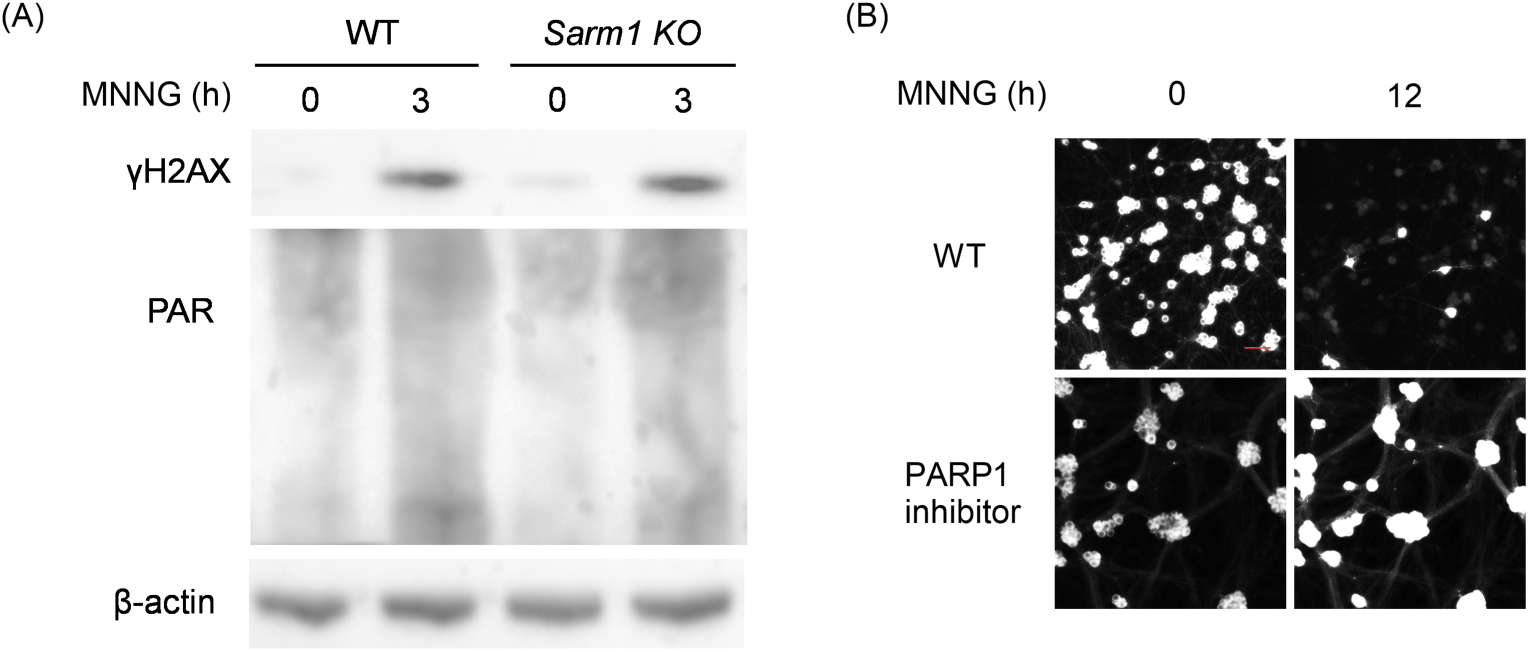
(A) Western blot analysis of γH2AX and PAR in WT and *Sarm1* KO DRG neurons treated with MNNG (500 µM) for 3 hr. (B) DRG neurons preincubated with PARP1 inhibitor (EB-47) were treated with MNNG (500 µM) at DIV7 and stained with TMRM dye to assess mitochondrial membrane potential. Scale bars, 100 µm.

**Supplemental Fig. 3:**
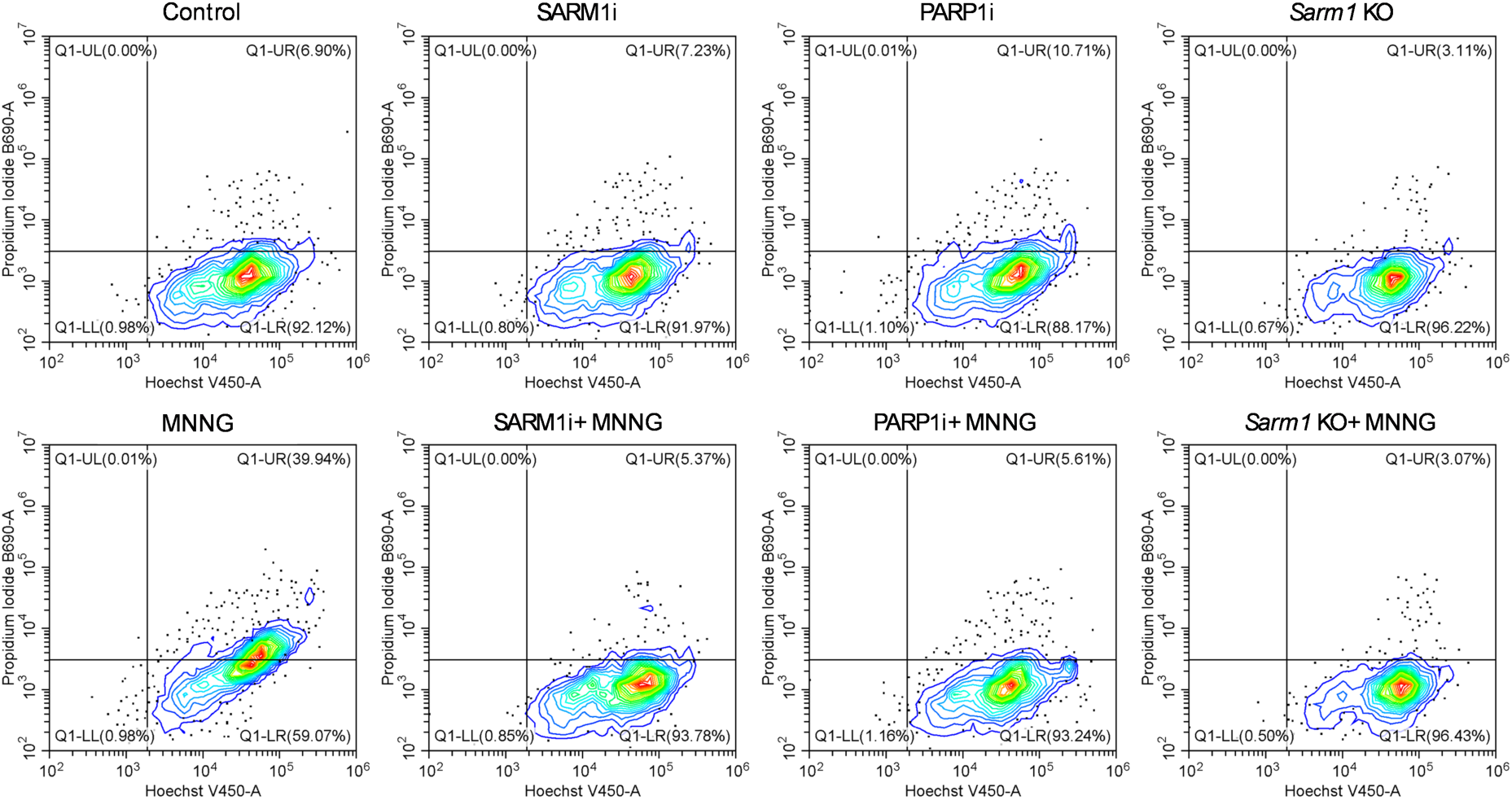
Flow cytometry to quantify cell death of WT or *Sarm1* KO DRG neurons preincubated with SARM1 inhibitor (NB-7) or PARP1 inhibitor (ABT-888) and treated with vehicle or MNNG (500 µM) for 12 hr. Cells were stained with Hoechst/PI. Upper right region shows percentage of dead cells after treatment.

**Supplemental Fig. 4:**
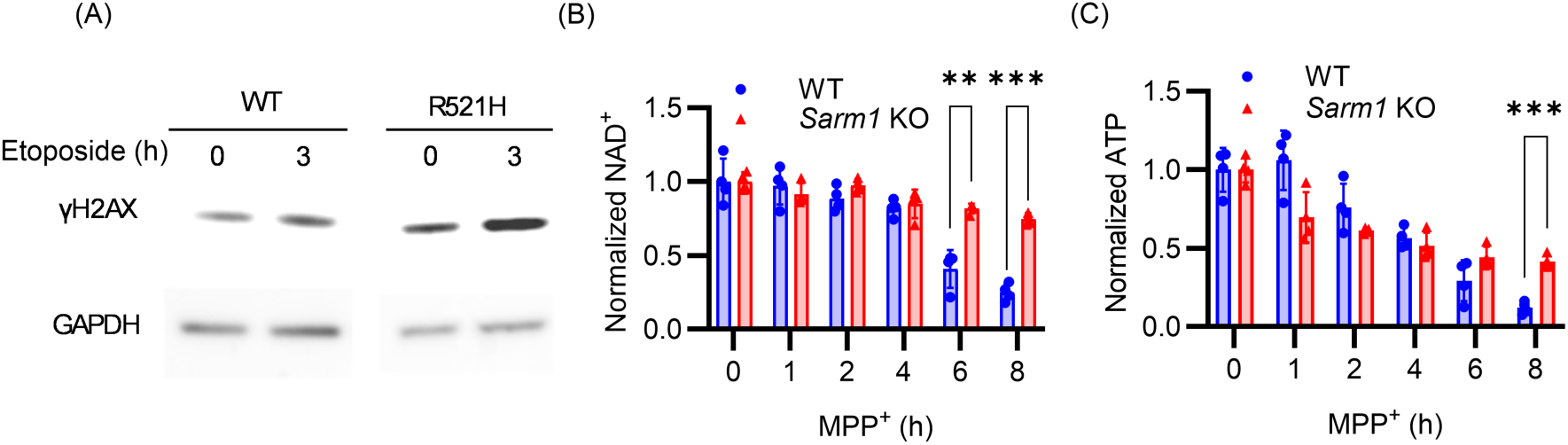
(A) Western blot analysis of the DNA damage marker γH2AX in *FUS^R521H^* and isogenic control (WT) iPSC-derived MNs treated with etoposide (10 µM) for 3 hr. (B,C) WT and *Sarm1* KO DRGs were treated with MPP^+^ (100 µM) at DIV7. NAD^+^ (B) and ATP (C) were quantified by LC-MS/MS at indicated times after treatment. Data with error bars correspond to Mean±SD. Statistical significance was determined by multiple unpaired t tests, comparing *Sarm1* KO to WT at each time point. **p < 0.01; ***P<0.001.

